# LiverMap pipeline for 3D imaging of human liver reveals volumetric spatial dysregulation of cirrhotic vasculobiliary architecture

**DOI:** 10.1101/2024.09.14.613049

**Authors:** Wesley B. Fabyan, Chelsea L. Fortin, Heidi L. Kenerson, Susana P. Simmonds, Jonathan T. C. Liu, Matthew M. Yeh, Rotonya M. Carr, Raymond S. W. Yeung, Kelly R. Stevens

## Abstract

The liver contains an intricate microstructure that is critical for liver function. Architectural disruption of this spatial structure is pathologic. Unfortunately, 2D histopathology – the gold standard for pathological understanding of many liver diseases – can misrepresent or leave gaps in our understanding of complex 3D structural features. Here, we utilized immunostaining, tissue clearing, microscopy, and computational software to create 3D multilobular reconstructions of both non-fibrotic and cirrhotic human liver tissue. We found that spatial architecture in human cirrhotic liver samples with varying etiologies had sinusoid zonation dysregulation, reduction in glutamine synthetase-expressing pericentral hepatocytes, regression of central vein networks, disruption of hepatic arterial networks, and fragmentation of biliary networks, which together suggest a pro-portalization/decentralization phenotype in cirrhotic tissue. Further implementation of 3D pathological analyses may provide a deeper understanding of cirrhotic pathobiology and inspire novel treatments for liver disease.

## Introduction

Microscopy stands as an indispensable tool in biological sciences, offering researchers the ability to delve into the intricate world of cells and tissues with exquisite detail. Since its inception in the 17^th^ century, microscopy has undergone numerous revolutionary advances propelled by innovations in optics, imaging technology, chemistries, and computational analyses [1]. Such innovations have led to vastly improved spatial resolution, imaging depth, and experimental throughput [1]. Recently, such advances have equipped scientists with a new toolkit to overcome limitations of microscopic imaging of 2D sections, which sliced convoluted structures into multiple objects, inaccurately captured object distribution, and routinely missed sparse objects [2]. This toolkit in tissue clearing, labeling, and imaging has enabled 3D volumetric imaging of tissues at cellular resolution, better elucidating tissue structure and its disease-associated alterations. Since tissue structure and function are intimately linked in biological tissues, such 3D analyses may provide a deeper understanding of functional abnormalities and inspire novel treatments for disease.

The liver is an exemplary organ where its 3D microarchitecture is integral to its proper function, and disruption to its structural architecture is pathologic [3]. Healthy liver tissue consists of structural units called lobules, which are thought to be polygonal in shape and tessellate throughout the liver. At the lobule vertices are the portal tracts, which contain portal veins, hepatic arterioles, and bile ducts amongst other structures, and at the lobule center is the central vein [4]. Blood enters the liver through portal tracts at the vertices of the lobule, percolates inwards through the parenchymal space via the liver sinusoids, and exits through the central vein at the center of the lobule [5] [6]. The lobule’s cellular organization is critical for its functional and structural zonation wherein cells are spatially divided into concentric zones that perform unique metabolic functions and/or exhibit unique structural features [7].

Liver cirrhosis – an advanced stage of liver disease marked by the increase of fibrotic scar tissue – may cause the liver’s intricately organized microvasculature to become massively rearranged and result in significant functional deficiencies. Cirrhosis typically originates from consistent exposure to damaging agents, such as alcohol (decompensated alcohol-associated cirrhosis), fatty acids (metabolic dysfunction-associated steatotic liver disease), hepatitis B or C viruses, and/or drug abuse or misuse [8]. In some cases, the root cause that leads to cirrhosis may be unknown, such as in primary sclerosing cholangitis which is a rare disease that results in cirrhosis characterized by bile duct inflammation and scarring [9]. The cirrhotic etiology may dictate the presentation of abnormal vascular and/or biliary architecture in a patient’s cirrhotic liver [10] [11] [12] [13]. Understanding how changes in liver architecture correlate with cirrhotic disease states and impaired liver function may provide avenues for understanding and potentially treating cirrhosis.

Despite the potential power of 3D imaging for shedding light on liver structure and pathology, most prior 3D imaging of the liver has been performed in rodent liver tissue [14] [15] [16] [17] [18] [19] [20] [21] [22] or in humans on the scale substantively less than a single liver lobule [23] [24]. For example, the largest prior human liver 3D imaging study with cellular resolution imaged from a portal vein to a central vein along the portal-central axis (∼500µm x 100µm x 100µm) [23]. To date, no published study has visualized non-fibrotic (*i.e.,* normal) or cirrhotic human liver tissue across multiple lobules with cellular resolution, which is key for understanding how the 3D lobular architecture of human liver tissue remodels during cirrhosis and results in functional deficiencies.

Here, we developed an accessible LiverMap pipeline to capture cellularly resolved human liver tissue on the mesoscale (4,000 x 4,000 x 500 µm), enabling visualization of several adjacent human lobules in 3D with cellular resolution over a depth of 500 µm. 3D comparative analyses of non-fibrotic control and cirrhotic tissues indicated that cirrhosis correlates with the dysregulation of sinusoid zonation, reduction in glutamine synthetase-expressing pericentral hepatocytes, regression of central vein networks, disruption of hepatic arterial networks, and fragmentation of biliary networks. Together, these observations suggest a pro-portalization/decentralization phenotype in correlation with human liver cirrhosis. The LiverMap pipeline thus offers an accessible method to gain a more holistic, empirical view of 3D human liver architecture and its cirrhosis-induced adaptive remodeling.

## Results

### LiverMap pipeline for 3D visualization of human liver tissue

To democratize the ability to visualize human liver tissue across several lobules with cellular resolution, we developed the LiverMap pipeline for 3D staining, clearing, reconstructing, and analyzing structures in the human liver. Towards this end, human non-fibrotic (*i.e.,* “normal liver”) control or cirrhotic liver samples were excised from either the livers of patients undergoing oncologic partial hepatectomy or from the explanted livers of patients receiving liver transplantation, respectively (Fig. 1a). Non-fibrotic control samples were obtained from resected non-diseased human liver tissue located at least two centimeters away from any present tumor, a distance at which tissue is typically considered free from field effect [25] [26]. For all human liver specimens used here, an adjacent slice was also sectioned and stained via traditional 2D histology methods and assigned an ISHAK score to denote the degree of fibrosis by a blinded pathologist (Suppl. Fig. 1). Next, each human liver sample was punched into 6mm discs and sliced to a thickness of 2mm (“liver slices”), a size expected to contain several human liver lobules (Fig. 1a). Samples were then immunostained with various antibodies to identify hepatocytes, liver sinusoidal endothelial cells (LSECs), portal and central veins, hepatic arterioles, bile ducts, and/or bile canaliculi (Table 1). Stained samples were optically cleared using the Ce3D protocol for lipid-rich, wholemount tissue (Fig. 1a) [27] and then tile-imaged with a confocal laser scanning microscope (Suppl. Fig. 2). Finally, commercially available software was utilized to computationally reconstruct immunofluorescent 3D human liver images via thresholding, segmentation, and binary interpretation for visualization and quantification (Fig. 1b,c; Suppl. Fig. 3; Suppl. Fig. 4; Suppl. Fig 5).

**Figure 1:**
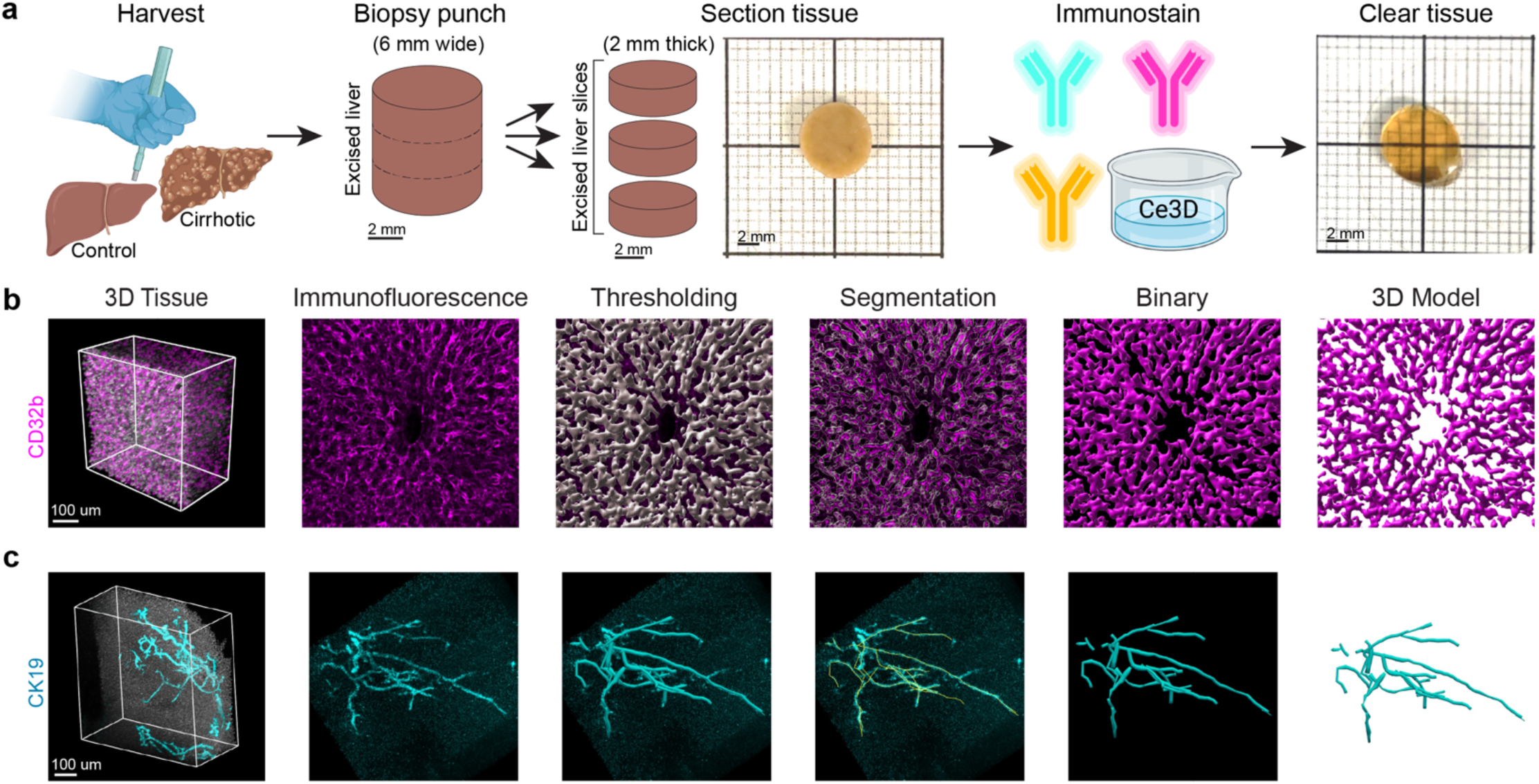
Schematic of the LiverMap pipeline. **a**, Human liver tissue is collected during surgical resection of non-fibrotic control liver (Harvest, left) and cirrhotic liver (Harvest, right). Next, biopsy punches are taken of the excised liver tissue and sliced into 2 mm thick sections before being stained with fluorescent molecules or antibodies and optically cleared via Ce3D for 3D imaging. **b**, **c**, After cleared tissues are imaged in 3D, immunofluorescent signals are used to computationally create object surfaces using Imaris 9.7 software (**b**) and vessel networks using Vesselucida software (**c**) for visualization and quantification.

**Table 1:**
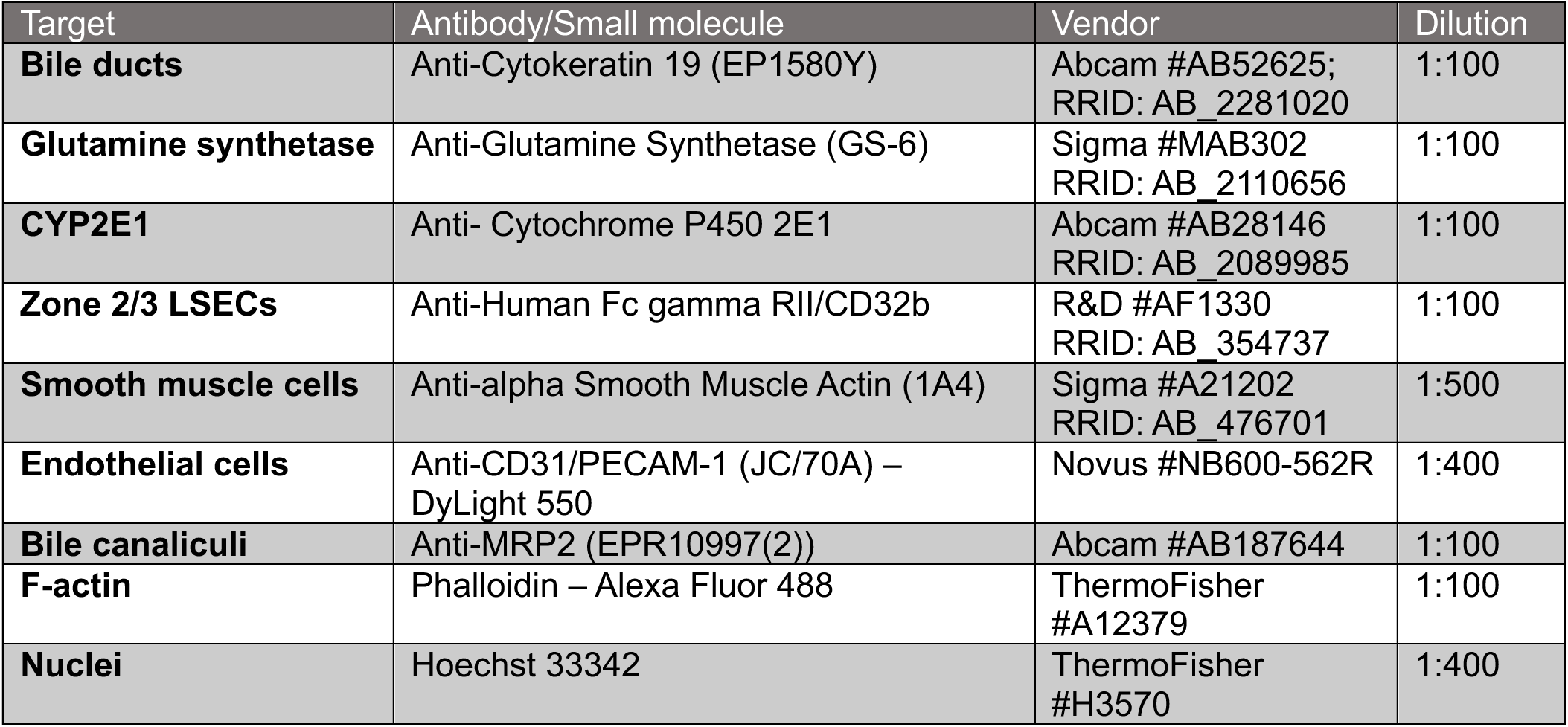
Antibodies and small molecules.

### Multiscale visualization of architectural features of human liver tissue

We leveraged the LiverMap pipeline to expand upon prior human liver tissue 3D imaging volumes to map how cellular and structural features were arranged across several length scales in 3D space. To do this, we first visualized how multiple lobules tessellate across a volume of human liver (Fig. 2a, left; Fig. 2b). We identified the center of each polygonal liver lobule, the central vein which transports processed blood out of the liver (Fig. 2a, center), by immunostaining the hepatocytes immediately surrounding the central vein using glutamine synthetase (GLUL; Fig. 2b) and manually tracing the central vein lumen through the 3D image stack. The polygonal vertices at the peripheral edge of each lobule were identified by immunostaining the bile ducts in portal tracts via cytokeratin 19 (CK19; Fig. 2b) [28] [29]. We were able to visualize and reconstruct portions of 17 different liver lobules across a 4mm x 4mm x 500µm volumetric space (Fig. 2b). The median average human liver lobule polygonal radius was 595µm (n=17; range: 428 to 717µm), which is consistent with the findings of prior studies of the human liver [23] [30] [31] [32].

**Figure 2.**
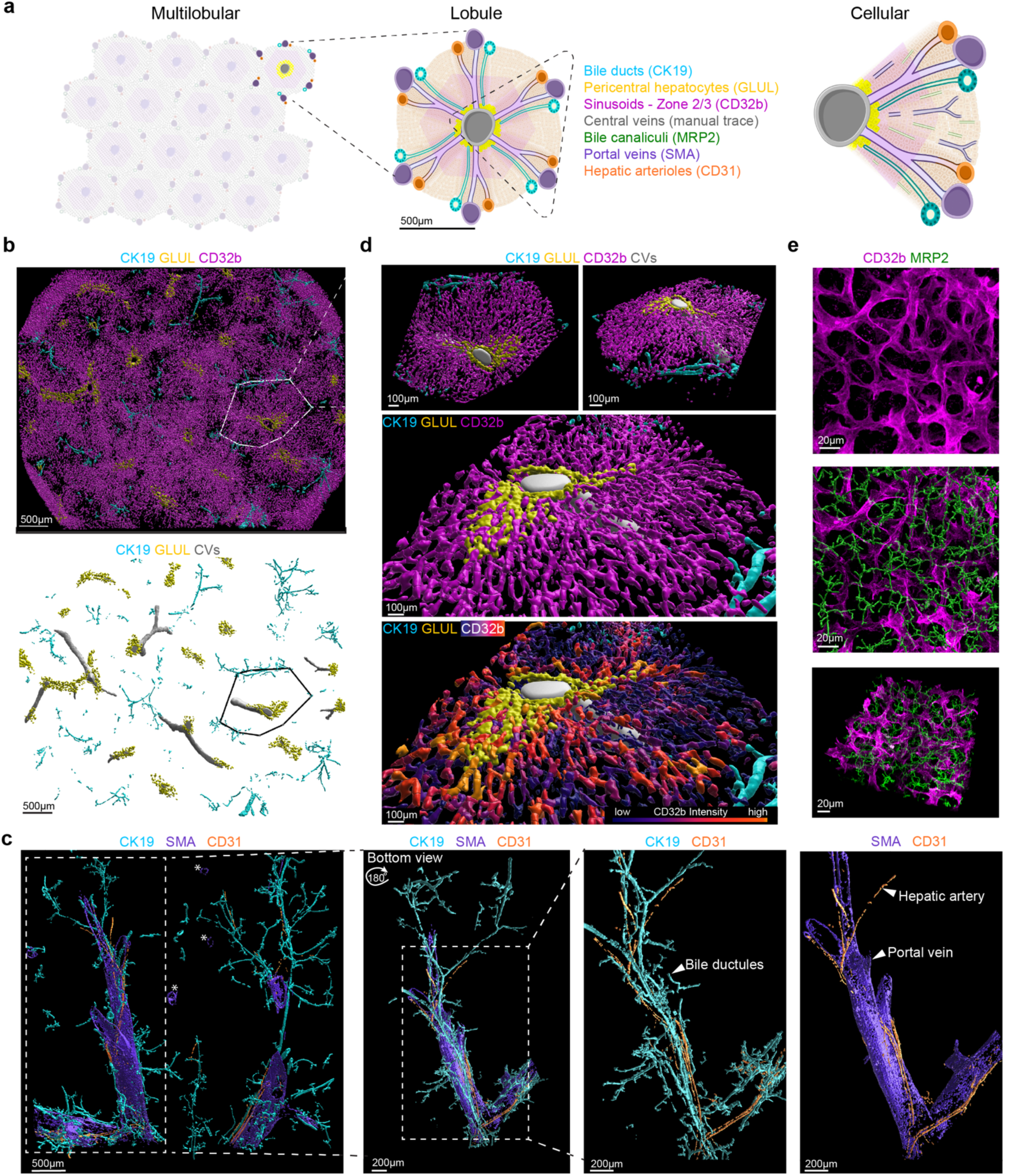
Multilobular reconstructions of zonated and architectural features. **a**, Native liver structure consists of repeating polygonal functional units called lobules, which have a central vein (SMA) at the center, Zone 2/3 LSECs (CD32) and pericentral hepatocytes (GLUL) radially organized around the central vein, and portal tracts (portal veins, SMA; bile ducts, CK19; hepatic arterioles, CD31) at the vertices. **b**, Multilobular imaging and 3D modeling of zonation features: Zone 2/3 LSECs, bile ducts, and pericentral hepatocytes. CD32b (magenta) is expressed by Zone 2/3 LSECs. CK19 (cyan) marks biliary epithelial cells. GLUL (yellow) labels pericentral hepatocytes. Top, Imaris surface rendering of CK19, GLUL, and CD32b immunofluorescent signals with an approximate lobule border (white polygon). Bottom, Imaris surface rendering of CK19 and GLUL immunofluorescent signals with manually traced central veins (grey) and an approximate lobule border (black polygon). **c**, Multilobular imaging and 3D reconstructions of architectural landmarks: portal veins, central veins, bile ducts and hepatic arterioles. SMA (purple) marks smooth muscle cells of portal veins and central veins; central veins are denoted by asterisks (left). CD31 (orange) labels vascular endothelial cells, especially bright on hepatic arterial endothelium. CK19 (cyan) marks biliary epithelial cells. Left, Whole-biopsy punch image of CK19, SMA, and CD31 surface renderings. Center-left, Bottom view of the zoom section in the left panel rotated 180 degrees about the y-axis. Center-right, Zoom-in of bile ducts (CK19) and hepatic arterioles (CD31). Right, Zoom-in of the larger portal vein (SMA) and wrapping hepatic arterioles (CD31). **d**, Top, Zoom-in of isolated lobule in **b** (left, top-view; right, angled-view). Middle, Close-up of lobule showing GLUL-expressing hepatocytes surrounding the central vein (grey). Bottom, Intensity-based pseudo-coloring of CD32b expression showing that the highest CD32b signal expression (orange) is located near the central vein and decreases with distance along the central-portal axis. **e**, Immunofluorescent image of Zone 2/3 LSECs (top, magenta, CD32b) with bile canaliculi (green, MRP2) surface renderings (center) or raw MRP2 immunofluorescent signal (bottom).

We next sought to map the volumetric topology of mesoscale transport networks in the portal tracts and lobules. To do this, we stained, imaged, and reconstructed portal veins that bring partly deoxygenated but nutrient-rich blood to the liver, hepatic arterioles that bring oxygenated blood into the liver, and bile ducts that carry the hepatic byproduct bile out of the liver to the gallbladder and the small intestine (SMA, CD31, and CK19 respectively; Fig. 2c). SMA also stains central veins and thus demarcated the center of lobules (Fig. 2c, SMA marked by asterisks). The resultant 3D visualizations demonstrated how bile ducts (CK19/cyan) and hepatic arterioles (CD31/orange) extend, wrap, encircle, and branch alongside large interlobular portal veins (SMA/purple) (Fig. 2c; Suppl. Video 1). These reconstructions volumetrically displayed how several portal tracts together encase a liver lobule (Fig. 2c, left, lobule center denoted by asterisks-labeled central vein).

As a next step, we sought to further visualize transport networks within a given lobule (Fig. 2a, center). We first used CD32b to visualize vascular sinusoids, which are specialized capillaries lined with highly permeable endothelial cells (i.e., LSECs) that facilitate macromolecular exchange between the blood and hepatocytes. 2D histological studies have previously shown this marker is highly expressed in Zone 2/3 human LSECs in the mid-lobular and pericentral regions of the lobule and lowly expressed in Zone 1 human LSECs in the portal regions of the lobule (Fig. 2a, center) [33]. To visualize the sinusoids, we computationally created binary reconstructions of immunofluorescent CD32b signal to visualize all cells expressing CD32b regardless of expression level (Fig. 2b,d, magenta). We then isolated a single lobule via Imaris clipping planes using portal features as lobular boundaries. LSEC zonation was further visualized by pseudo-coloring immunofluorescent signal intensity (Fig. 2d, bottom), which showed roughly a radial gradient pattern with LSECs most strongly expressing CD32b at the central region of the lobule.

We further sought to visualize not only microscale vascular transport networks within the lobule but also the bile canalicular networks, which are 1-2µm wide interconnected spaces between adjacent hepatocytes that collect and carry byproduct bile salts produced by hepatocytes to the bile ductules [4]. To do this, we immunostained for multidrug resistance-associated protein (MRP2), a transporter located on the canalicular (apical) surface of hepatocytes [34]. The resultant cellular and network-level volumetric images of CD32b^+^ LSECs and MRP2^+^ bile canaliculi depicted the complexity of how these two micro-transport networks interweave in 3D (Fig. 2e). These surfaces also demarcated hepatocyte location, as these cells have surfaces that face both the sinusoids (Fig. 2e, magenta) and bile canaliculi (Fig. 2e, green). Altogether, the LiverMap pipeline enabled capture and 3D visualization of cellularly resolved tissue volumes that are ∼1,600-fold larger than prior studies for multiscale reconstruction of 3D human liver structure.

### Dysregulation of human hepatocyte and sinusoid zonation in cirrhosis

Zonation of the liver lobule is essential for healthy liver function as it allows the numerous tasks of the liver to be delegated across cell subpopulations [35]. Most prior investigations of liver zonation have focused primarily on hepatocytes, such as defining hepatocyte periportal functions and pericentral functions. Interestingly, recent research suggests that LSECs are also spatially zonated across the lobule [33]. However, these studies relied on 2D traditional methods and characterized only LSECs in normal liver tissue. The extent to which LSECs are volumetrically zonated in both normal and cirrhotic tissue remains unclear.

To address this knowledge gap, we first sought to benchmark how LSEC zonation correlates with hepatocyte zonation [35]. To do this, we employed the LiverMap pipeline to visualize the extent to which LSEC expression of CD32b correlates spatially with hepatocyte expression of zonation markers cytochrome P450 2E1 and glutamine synthetase (CYP2E1; GLUL; Fig. 3). We found that expression levels of CD32b in LSECs and CYP2E1 in hepatocytes in normal non-fibrotic human liver tissue closely correlated spatially across both multilobular line profiles and 3D signal intensity plots, exhibiting gradient patterns that emanated from the center of the lobules (Fig. 3a-c). Conversely, GLUL was confined to the innermost layers of hepatocytes surrounding the central vein, as shown previously by us and others [36], and thus overlapped spatially with only the most centrally located CD32b^+^ LSECs (Fig. 2; Fig. 3d,e).

**Figure 3:**
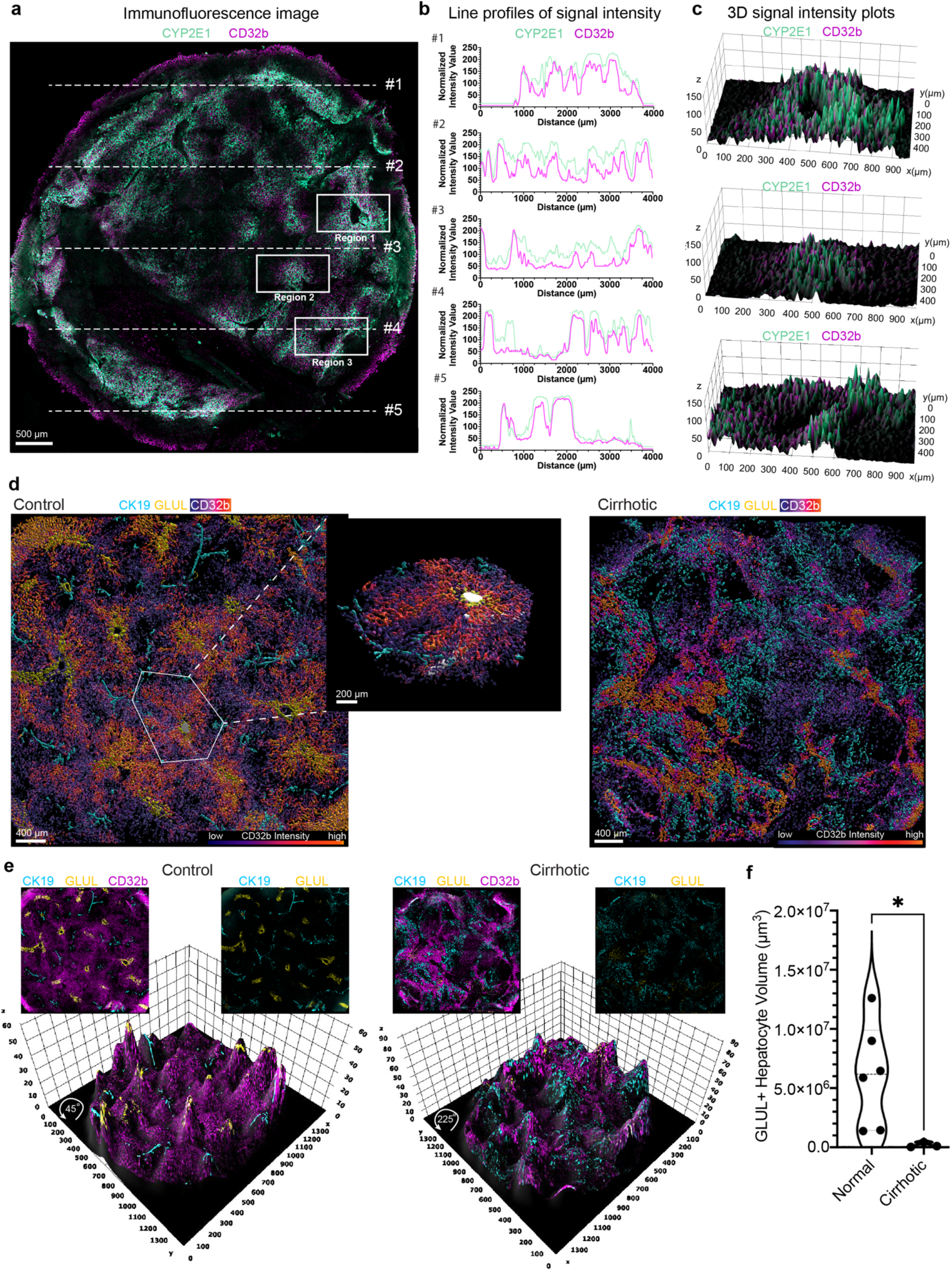
CD32b+ LSECs associate with CYP2E1+ hepatocytes, and 3D modeling reveals dysregulation of human sinusoid zonation in cirrhosis. **a**, En face 3D MIP of control tissue with CD32b (magenta) staining Zone 2/3 LSECs and CYP2E1 (teal) staining metabolically active pericentral hepatocytes. **b**, Spatial line profiles of CD32b signal intensity (magenta) and CYP2E1 signal intensity (teal) across horizontal lines denoted in **a**. From these line profiles, we can see that CD32b and CYP2E1 signals follow similar spatial trends across the sample. **c**, Angled 3D signal intensity plots showing pixel intensity on the z-axis corresponding to each pixel on the xy-plane of the three regions denoted in **a**. **d**, Left, control sample with CD32b surface pseudo-colored based on signal intensity. CD32b-high regions (orange) are typically in the central and midlobule zones, and CD32b-low regions (blue) are portal. Approximate lobule border drawn in white with inset showing an angled view of the isolated lobule from control tissue with a manually traced central vein. Right, Cirrhotic sample with CD32b surface pseudo-colored based on signal intensity. CD32b-high regions are less regularly patterned than seen in the non-fibrotic control sample. **e**, Top, En face MIP of entire non-fibrotic control (left) and cirrhotic (right) biopsy punches labeled for CK19+ bile ducts (cyan), GLUL+ pericentral hepatocytes (yellow), and CD32b+ Zone 2/3 LSECs (magenta). Bottom, Angled 3D signal intensity plots showing pixel intensity (z-axis) corresponding to each pixel of the 2D image (xy-plane) for CK19, GLUL, and CD32b. **f**, Statistical comparison of GLUL+ hepatocytes Imaris surface volumes for non-fibrotic control (n=6) and cirrhotic (n=3) tissues using a two-tailed Mann-Whitney test (*, p<0.05; p-value = 0.0238).

As the microvasculature can become disrupted during cirrhosis, we next sought to visualize spatial zonation patterns of LSECs across several lobules in both non-fibrotic and cirrhotic tissue. We stained six non-fibrotic and three cirrhotic liver slices to identify sinusoidal endothelial cells, bile ducts, and pericentral hepatocytes (CD32b, CK19, GLUL respectively; Suppl. Fig. 2a) [37]. We then created intensity-based pseudo-colored plots of CD32b immunofluorescent signal intensity across the multilobular tissue volume (Fig. 3d, orange denotes the highest signal & dark blue represents the lowest signal) as well as 3D surface plots of CD32b, CK19, and GLUL signal intensity from MIP images (Fig. 3e, z-values represent signal intensity). In non-fibrotic control human liver tissue, CD32b was most highly expressed at the center of each liver lobule, nearest to GLUL-expressing hepatocytes, and displayed roughly radial expression gradients emanating from the pericentral region of each lobule and abating at the lobule edges, demarcated by CK19^+^ bile ducts (Fig. 3d,e, left). Each roughly radial gradient of CD32b expression associated with each given liver lobule appeared to be interspersed regularly across non-fibrotic control human liver tissue. Conversely, in the cirrhotic tissue, this regular pattern of radial CD32b expression gradients was disrupted, with some but not all cells retaining CD32b expression (Fig. 3d,e, right). Furthermore, CD32b expression appeared to be interlaced with CK19 expression, suggesting that spatial dysregulation of zonated CD32b expression may accompany lobular vasculobiliary architectural changes in the cirrhotic liver tissue. Lastly, similarly to that shown by others previously in 2D sections [38], we found a near absence of volumetric GLUL expression in the cirrhotic samples (Fig. 3f; Suppl. Fig. 4; Suppl. Fig. 5), which may imply that CD32b dysregulation could be linked to disruption of hepatic zonation patterns and/or hepatocellular function.

In summary, the LiverMap pipeline enables visualization of volumetric functional and structural sinusoidal and hepatocyte zonation across several human liver lobules in 3D, and through leveraging this pipeline, we found that cirrhosis correlates with the dysregulation of sinusoid zonation and a reduction in glutamine synthetase-expressing pericentral hepatocytes.

### Alterations to liver vascular networks in cirrhotic tissue

Functional zonation is largely associated with the vasculobiliary structural hallmarks of the lobules, such as the central veins from which zonation patterns typically emanate. We and others have reasoned that disruption in zonation patterns, such as the reduction we observed in GLUL-expressing hepatocytes, may stem from disrupted central vein network architecture [11] [39]. We thus next set out to further map mesoscale 3D vasculobiliary network architectures in both non-fibrotic and cirrhotic human liver tissue.

To do this, we first reconstructed the central vein, hepatic arterial, and biliary networks within six non-fibrotic control tissues and three cirrhotic tissues (Fig. 4a,b). Parameterization and quantification of central vein networks demonstrated that normal non-fibrotic liver tissues had a higher number of central veins, a greater total network volume, and more network endings than cirrhotic tissues (Fig. 4c). While all cirrhotic tissue samples exhibited a reduction in central veins compared to normal non-fibrotic tissue, each cirrhotic sample had distinct architectural arrangements of mesoscale hepatic arterial networks (Fig. 4b). Specifically, we observed an expansion of the CD31^+^ vessels demarcating hepatic arterioles and possibly dysregulated sinusoids, which may become CD31^+^ after sinusoidal defenestration or capillarization [40] [41], in the samples from patients with decompensated alcohol-associated cirrhosis and primary sclerosing cholangitis (Fig. 4b, top and bottom, respectively) but not in the sample from the patient with hepatitis B viral infection (Fig. 4b, center). The different presentations of vascular architectures within the cirrhotic group may be attributed to samples having different cirrhotic etiologies or to differences in disease progression across patients. Altogether, these volumetric findings indicated and allowed the visualization of central vein network regression and hepatic arterial network disruption in cirrhotic human liver tissue.

**Figure 4.**
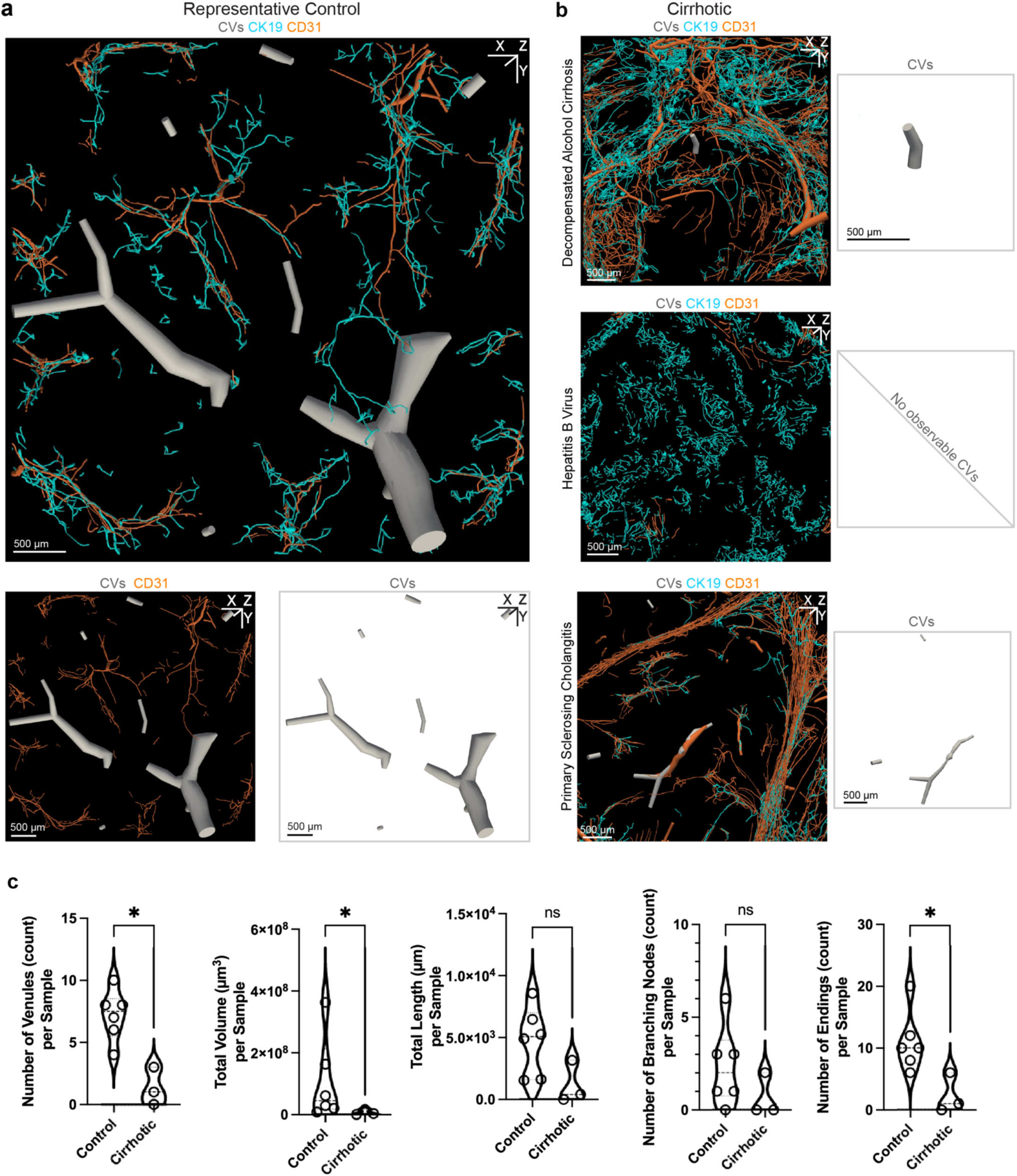
3D reconstructions reveal a decentralization of blood vessels in cirrhosis. **a**, Top, En face 3D reconstructions of identifiable central veins (grey), CK19+ bile ducts (cyan), and CD31+ microvessels (orange) across representative non-fibrotic control human liver tissue. Bottom, depiction of vasculature (grey and orange) isolated from non-fibrotic control 3D model above. **b**, Left, En face 3D models of identifiable central veins (grey), CK19+ bile ducts (cyan), and CD31+ microvessels (orange) across three cirrhotic human liver tissue sections with different etiologies. Right, 3D models of identifiable central veins (grey) visually isolated from adjacent cirrhotic samples. **c**, Morphometric measurements describing each sample’s central vein network in the control (n=6) and cirrhotic (n=3) groups. Data are represented by violin plots and were statistically analyzed via a two-tailed Mann-Whitney test (*, p<0.05; ns, non-significant).

### Architectural changes of the biliary network in cirrhosis

Bile ducts carry hepatic bile out of the liver to the gallbladder and the small intestine, which helps facilitate the excretion of hepatic metabolites and intestinal absorption of lipids and fat-soluble vitamins [42]. In liver cirrhosis, biliary networks can undergo significant remodeling and fail to transport bile out of the liver, thereby causing a buildup of bile in the liver and prompting bile leakage into the bloodstream – a condition called cholestasis [43]. Our initial vasculobiliary network tracings suggested architectural dysregulation in the CK19 biliary networks from patients with cirrhosis (Fig. 4b, cyan). We thus lastly sought to further investigate bile duct architecture in both normal and cirrhotic human liver tissue.

To do this, we created Imaris surface renderings of CK19 immunostains within six non-fibrotic samples and three cirrhotic samples (Fig. 5; Suppl. Fig. 5). In the non-fibrotic group, we observed biliary structures that extended and branched through tissue volumes, resembling a network architecture (Fig. 5a; Suppl. Fig. 5). However, in the cirrhotic group, each cirrhotic sample presented a unique CK19 biliary network architecture, which may be a result of the cirrhotic etiology or disease severity within each sample (Fig. 5b; Suppl. Fig. 5). We observed ductular expansion primarily within the thick fibrotic bands in the decompensated alcohol-associated cirrhosis sample, discrete clusters of CK19^+^ cellular structures throughout the hepatitis B cirrhosis sample, and a combination of long bile ducts within thin fibrotic bands and bile duct segments within the bulk of the tissue in the primary sclerosing cholangitis sample (Fig. 5b; Suppl. Fig. 2; Suppl. Fig. 5).

**Figure 5:**
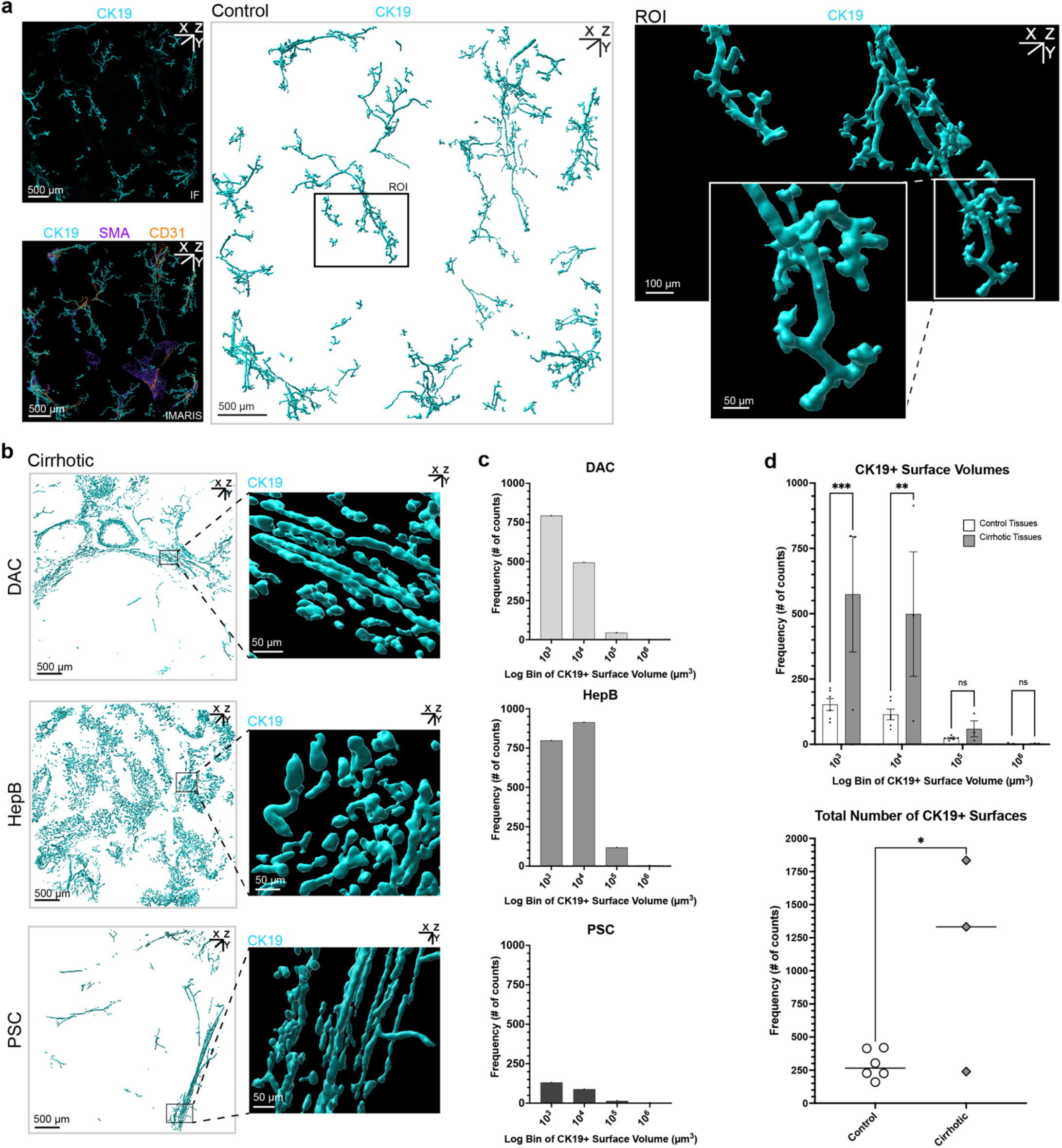
Bile duct network consists of more architectural fragmentation in cirrhosis. **a**, Top left, En face immunofluorescence MIP of CK19^+^ bile ducts in a representative non-fibrotic control sample. Bottom left, Imaris surfaces of CK19^+^ bile ducts, SMA^+^ portal veins, and CD31^+^ hepatic arterioles from immunofluorescence signal in representative non-fibrotic control sample. Center, Imaris surfaces of CK19^+^ bile ducts in representative control sample with ROI box. Right, Zoom-in of ROI identified in the center panel with high zoom-in panel showing a selected bile duct. **b**, Imaris surfaces of CK19^+^ features in the cirrhotic group with different etiologies (DAC, decompensated alcohol-associated cirrhosis; HepB, hepatitis B virus; PSC, primary sclerosing cholangitis). **c**, CK19 Imaris surface volumes from cirrhotic samples binned with regard to their log base 10 to create exponential histograms of CK19 surface volumes for each sample. **d**, Top, Comparison of non-fibrotic control (n=6) and cirrhotic (n=3) CK19 surface volume exponential histograms (error bars, +/-SEM). Exponential histogram bins were statistically compared via two-way ANOVA (F = 14.90; DFn = 1; DFd = 28) and Bonferroni multiple comparison test (***, p<0.001; **, p<0.01; ns, non-significant). Bottom, Comparison of the total number of CK19 surfaces within non-fibrotic control (n=6) and cirrhotic (n=3) groups. Average total number of CK19 surfaces between groups were statistically compared via a two-tailed, unpaired t-test (t = 2.679; df = 7; *, p<0.05).

For the cirrhotic samples, we were particularly interested in 3D morphological analysis of a pathological bile duct phenotype called ductular reaction (DR), which resembles an expansion of small bile ductules and may be caused by the proliferation of cholangiocytes or hepatic progenitor cells, or by transdifferentiation of hepatocytes [12]. DR is not fully understood, but it is believed that the cell type of origin for DR may depend on the underlying pathology that caused the cirrhosis [12] [13]. Typically, 2D histology is used for the evaluation of ductular reaction; however, 2D analysis often cannot elucidate the extent to which such 3D structures are interconnected within the tissue, especially when structures are lumenized. As examples, we point to the morphological similarity of CK19^+^ features in several 5µm-thick slices across two different human liver specimens (Suppl. Fig. 6). Upon 2D inspection, the two samples appeared to have similar presentation of CK19^+^ features (Suppl. Fig. 6, left). However, using 3D volumetric imaging, we observed that one sample had CK19^+^ bile ducts forming interconnected networks (Suppl. Fig. 6, top right), and the other sample had CK19^+^ features representing distinct, non-connected smaller structures (Suppl. Fig. 6, bottom right). To our knowledge, ductular reaction architecture has not yet been quantifiably measured across several human liver lobules in 3D.

Towards this end, we calculated the volume of each discrete bile ductular structure in every non-fibrotic control and cirrhotic sample (Suppl. Fig. 4; Suppl. Fig. 5) and generated exponential histograms of CK19^+^ ductular volumes for each sample (Fig. 5c). This method provides a new quantitative means to describe differences in ductular fragmentation between different diseased samples, a metric that is not possible to accurately derive from 2D images (Fig. 5c; Suppl. Fig. 6). Here, the three samples with different cirrhotic etiologies had histograms of markedly different shapes (Fig. 5c), which reflected their distinctive ductular architectures visualized in 3D (Fig. 5b). The frequency counts for non-fibrotic or cirrhotic samples were then grouped and plotted as average frequency distributions (Fig. 5d, top). The resultant analyses showed that overall, cirrhotic samples on average had both a higher quantity of the smallest (<10^3^ um^3^ and <10^4^ um^3^) CK19 structures (Fig. 5d, top) and a higher total number of CK19 structures (Fig. 5d, bottom), compared to non-fibrotic samples. Altogether, our method thus provided a means to quantitatively describe ductular remodeling and fragmentation in human liver cirrhosis, enabling us to quantitatively visualize the differing degrees of ductular fragmentation regarding each cirrhotic sample as well as conclude that, on average, cirrhotic samples had more fragmented ductular networks than those in the non-fibrotic human liver control samples.

## Discussion

In this work, we utilized immunostaining, tissue clearing, 3D imaging, and computational modeling to develop an accessible LiverMap pipeline (Fig. 1). The resultant 3D images had a depth equivalent to ∼100 traditional tissue sections and a volume roughly 1,600 times larger than the previously captured human liver volume at cellular resolution (Fig. 2; Suppl. Fig. 2). These efforts unveiled never-before-seen-in-3D human liver structures across multiple size scales and visualized volumetric dysregulations of vasculobiliary architectures in cirrhosis.

Interestingly, while we did find shared vasculobiliary architectural changes across all cirrhotic samples compared to non-cirrhotic tissue (Fig. 4c; Fig. 5d), we also identified distinct vasculobiliary architectural features across cirrhotic samples (Fig. 4b; Fig. 5b,c). Further studies are thus needed to parse out key differences in vasculobiliary presentation across individuals with cirrhosis. While numerous factors could contribute to architectural variation across individuals with cirrhosis, two possibilities are the degree of cirrhosis, which was similar across our cirrhotic samples as per ISHAK scoring (Suppl. Fig. 2c), and cirrhotic etiology [10] [11] [12] [13], which importantly was different for each cirrhotic sample here. Notably, the LiverMap pipeline now provides a tool to perform such future studies.

Through leveraging the LiverMap pipeline, we observed a spatial dysregulation of CD32b^+^ LSECs in cirrhotic tissue in comparison to non-fibrotic control tissue (Fig. 3), which could imply a disruption in critical Zone 2/3 LSECs roles such as functions related to blood-derived leukocyte adhesion and antigen clearance in cirrhotic tissue [33]. Additionally, we found a significant reduction in glutamine synthetase-expressing pericentral hepatocytes in cirrhotic tissue when compared to non-fibrotic tissue (Fig. 3f). As glutamine synthetase plays an important enzymatic role in nitrogen metabolism by catalyzing glutamate and ammonia to form glutamine, this observation may imply decreased ammonia metabolism for cirrhotic patients, thereby indicating a higher risk of hyperammonemia or a buildup of toxic ammonia in the blood [44] as is commonly observed in patients with cirrhosis and liver failure [44] [45]. Furthermore, our 3D analyses revealed a significant decrease in the presence of central veins within cirrhotic tissue in comparison to non-fibrotic tissue (Fig. 4). Since the central vein serves as the primary transporter of blood out of the lobule, fewer central veins may result in dysfunctional blood drainage from the liver and potentially contribute to portal vein hypertension, a clinically observed condition within cirrhotic patients [10] [11] [46].

Lastly, we provided a new quantitative means to describe CK19 biliary network fragmentation in non-fibrotic and cirrhotic human tissues. We found that the presentation of CK19 biliary networks varied substantially between cirrhotic etiologies (Fig. 5; Suppl. Fig. 5), which was reflected by each cirrhotic sample having CK19 volumetric histograms with markedly different distributions (Fig. 5c). We also found that, on average, cirrhotic samples had both a higher quantity of the smallest CK19 structures (Fig. 5d, top) and a higher total number of CK19 structures (Fig. 5d, bottom) than non-fibrotic samples. Together, these statistics indicated that the average cirrhotic bile ductular network was more fragmented than the average non-fibrotic control bile ductular network, which may imply abnormal biliary pressure and flow in cirrhotic tissue.

In summary, this work introduced the LiverMap pipeline and presented the first 3D visualization and volumetric characterization of sparse and convoluted structures across several lobules in human liver tissue. With this novel capability, we examined both non-fibrotic and cirrhotic human liver tissue to identify significant functional and architectural abnormalities associated with cirrhosis. Our 3D comparative analyses of non-fibrotic control and cirrhotic tissues indicated that cirrhosis correlates with the dysregulation of sinusoid zonation, reduction in glutamine synthetase-expressing pericentral hepatocytes, regression of central vein networks, disruption of hepatic arterial networks, and fragmentation of biliary networks. Together, these observations suggest a pro-portalization/decentralization phenotype in correlation with human liver cirrhosis. As liver structure is closely related to its function, further implementation of the LiverMap pipeline may enhance our 3D pathological understanding of architectural remodeling and functional deficiencies during cirrhosis and inspire novel treatments for liver disease.

## Methods

### Human liver samples

Patients undergoing surgical resection at the University of Washington Medical Center consented under an IRB-approved protocol (#00001852, Liver and Abdominal Tumor Biorepository) for tissue donation and access to de-identified demographic and clinical data. Non-cirrhotic control samples were obtained from partial hepatectomy (n=6), where each sample was dissected at least two centimeters away from any present tumor to ensure the sample was free from mass effect [24] [25]. Cirrhotic liver samples were collected from patients undergoing transplant (n=3). Tissue sections (∼2 x 2 x 2cm) not needed for pathologic evaluation were fixed in 10% formalin and sliced with a vibratome (Leica VT 1200S) to a thickness of 2mm. Tissue samples were fixed for 48 to 72 hours at 4°C before being transferred to phosphate-buffered saline at 4°C until further use. Pieces of each tissue sample were sectioned and stained with hematoxylin and eosin (H&E) and Masson trichrome (MT) to evaluate tissue structure (H&E) and fibrosis content (MT) using conventional methods (Suppl. Fig. 2). A pathologist then performed a blinded ISHAK grading of the H&E and MT images for each patient sample to confirm the non-fibrotic and cirrhotic tissue groupings (Suppl. Fig. 2c). The histological, demographic, pathologic, and clinical details of all samples in this work are summarized in Supplemental Figure 1.

### LiverMap immunostaining and tissue clearing

For immunostaining, tissue samples were stained with antibodies or small molecules to identify bile ducts (cytokeratin 19, CK19), pericentral hepatocytes (glutamine synthetase, GLUL; cytochrome P450 2E1, CYP2E1), Zone 2/3 LSECs (CD32b), portal and central veins (smooth muscle actin, SMA), hepatic arterioles (CD31), bile canaliculi (multidrug resistance-associated protein 2, MRP2), F-actin filaments (phalloidin) and/or cell nuclei (Hoechst) (Table 1). For tissue clearing, the Ce3D protocol for lipid-rich, wholemount tissue was utilized [27].

Briefly, human liver tissue samples were punched into 6 mm discs using biopsy punches, sliced to a thickness of 2mm, washed in fresh phosphate-buffered saline (PBS) for one hour, and then blocked in Ce3D alternative blocking buffer (1x BD Perm/Wash buffer + 1% bovine serum albumin + 1% normal donkey serum + 0.3% Triton X-100 in PBS) overnight at 37°C with gentle shaking. Next, samples were incubated in primary antibodies diluted in Ce3D alternative blocking buffer with 5% dimethyl sulfoxide (DMSO) for 24 hours at 37°C with gentle shaking (Table 1). If primary antibodies were conjugated to a fluorophore, samples were protected from light from this step on. Samples were then washed with Ce3D wash buffer (0.2% Triton X-100 + 0.5% 1-thioglycerol in PBS) for six to eight hours at room temperature with gentle shaking, changing the buffer every two hours. Washed samples were then incubated in fluorophore-conjugated secondary antibodies diluted 1:400 in Ce3D alternative blocking buffer with 5% DMSO overnight at 37°C with gentle shaking and were protected from light from this step on. Samples were next washed in Ce3D wash buffer for six to eight hours at room temperature with gentle shaking, changing the buffer every two hours. Finally, samples were transferred to Ce3D clearing solution with Hoechst 33342 (1:400 dilution) for overnight incubation at 37°C with gentle shaking, allowing for water-clearing solution exchange within the tissue. After 12 hours, the samples were transferred to fresh Ce3D clearing solution and imaged. For long-term storage, samples were placed in multi-well plates, and plates were covered in parafilm and aluminum foil and stored at 4°C.

### Confocal imaging and image processing

Human liver samples were submerged in Ce3D clearing solution within glass-bottom dishes and covered with coverslips to keep the samples flat before being imaged on a confocal laser scanning microscope (Leica TCS SP8) with a 10x 0.4 numerical aperture dry objective (Leica). Hoechst, Alexa Fluor 488, Alexa Fluor 555 or DyLight 550, and Alexa Fluor 647 were excited with 405, 488, 552, and 638 laser lines, respectively, and detected with hybrid detectors (HyD). Z-stacks were acquired through Leica Application Suite X (LAS X) software with the bottom of the stack set ∼5 µm below the start of the visible signal and with the top of the stack set at the end of the visible signal, with a Z-compensation set within LAS X to increase laser power and gain throughout the stack as necessary. To capture the 6 mm biopsy punches, tiles of 4 x 4 image stacks were stitched using the Mosaic Merge function within LAS X. Stitched confocal Z-stack images were then visualized and processed using Imaris 9.7 (Bitplane). First, Z-stack images were converted from .lif file format to .ims using Imaris File Converter 9.7 (Bitplane). Images were then processed with the following steps: 1) Flip Channels in the Z direction for all channels; 2) Normalize Layers of all channels; 3) Background Subtraction (Filter Width: 10um) of all channels; 4) Normalize Layers of all channels, 5) Rotate Channels counter clockwise for all channels about X-axis, Normalize Layers for all channels, Rotate Channels clockwise for all channels about X-axis; 6) Rotate Channels counter clockwise for all channels about the Y-axis, Normalize Layers for all channels, Rotate Channels clockwise for all channels about Y-axis. All immunofluorescent images from this study after stitching and processing are shown in Supplemental Figure 2.

### Computational reconstructions for visualization and quantification

Imaris 9.7 (Bitplane) software was used to generate a surface rendering of the immunofluorescent signal from each antibody in all samples for enhanced visualization and feature volume quantification. The following parameters were used for the Imaris Surface Creation Wizard: Surface Grain Size = 4.01µm; Background Subtraction - Diameter of Largest Sphere = 15.0µm; Threshold - Background Subtraction manually adjusted for each sample to help account for signal variability; Voxel Filter or Volume Filter to exclude small surfaces created from background signal. Any significant background signal or tissue autofluorescence that was captured in the surface reconstructions was manually identified and removed. **Central vein visualization**: For visualization purposes, central veins were identified via positive glutamine synthetase staining or positive SMA staining outside of portal tracts, and central vein surfaces were created by manually tracing lumens throughout the tissue using autofluorescence (Fig. 2; Fig. 3). **Lobule measurements:** Lobule boundaries were identified and isolated via recognition of portal tracts and the use of Imaris clipping planes.

The Imaris measurement tool was then used to measure the distances between each central vein and all its nearest surrounding portal tracts for every full or partial lobule (n=17) identified in the representative sample (non-fibrotic #6). Each full or partial lobule’s measurements were then averaged together to get an average lobule radius for each full or partial lobule. Average lobule radii were then put in rank order, and the median lobule radius was identified and reported. **Immunofluorescent pseudo-coloring**: For intensity-based pseudo-coloring of the CD32b signal, the Split Touching Objects option was enabled and a custom color gradient with an Intensity Mean range of 3,000 units was utilized to visualize reconstructed sinusoid CD32b-expressing segments throughout the tissue sample. These Imaris surfaces aided in CD32b immunofluorescent signal visualization (Fig. 2d; Fig. 3d). **Glutamine synthetase volumetric measurements**: Each sample was processed to normalize the total image volume to 4,000 x 4,000 x 300 µm (Suppl. Fig. 4). For each sample’s normalized tissue volume, glutamine synthetase immunofluorescent signal was rendered into Imaris surfaces, and the total surface volume for each sample was calculated via the Imaris statistics module (Fig. 3f).

### Line profile analysis and 3D surface plots

2D maximum intensity projection (MIP) images of 500µm-thick z-stack images were exported from Imaris 9.7 software as .tiff files. The 2D MIP images were then opened in FIJI software [47]. Line profile analysis (Fig. 3a,b) was performed in Fiji with the following pipeline: 1) Convert the stack to 8-bit; 2) Apply a 15µm Gaussian Blur to help smooth the image and reduce noise; 3) Apply Equalize Histogram function to help normalize pixel intensities across images; 4) Employ Fiji’s Plot Profile analysis tool for equally spaced horizontal lines throughout the sample. 3D signal intensity plots (Fig. 3c,e) were also generated in FIJI by selecting a rectangular region of interest and employing Fiji’s 3D Surface Plot analysis tool with the following parameters: Grid Size: 512; Smoothing: 6.0; Perspective: 0.22; Lighting: 0.2; Scale: 1.73; z-Scale: 0.52.

### Central vein morphometrics

Vesselucida (MBF Bioscience) was used for the reconstruction and morphometric analysis of central veins (Fig. 4). Before performing morphometric analysis within Vesselucida software, images were resized to 25% using the Resize tool within LAS X software. This provided sufficient image resolution for analysis while reducing the file size to meet program and computing limitations. After resizing, a semi-manual process was used to trace branching structures in Vesselucida. First, the automatic tracing module was used to identify the microstructures with diameters of approximately 15 µm, such as CD31^+^ microvessels and small bile ductules. Then, larger vessels or ducts were manually traced, and these were manually connected to smaller automatically traced vessels. Central veins were manually traced in their entirety as these features were all too large for the automatic tracing in Vesselucida to identify reliably. From these tracings, Vesselucida generated morphometric data describing the architecture of traced networks.

### Ductular fragmentation quantification

The volume and quantity of each discrete CK19^+^ structure in every non-fibrotic control and cirrhotic sample was calculated and recorded via the Imaris statistics module (Suppl. Fig. 4; Suppl. Fig. 5). Exponential histograms of CK19^+^ surface volumes (bin width: log_10_ of surface volume in µm^3^) for each non-fibrotic control sample and cirrhotic sample were then created (Fig. 5c). The frequency counts of the resulting exponential histograms from each sample were then grouped with their respective non-fibrotic or cirrhotic classification and plotted as average frequency distributions (Fig. 5d, top), and frequency counts of CK19^+^ surface volumes within each bin from the non-fibrotic group and cirrhotic group were statistically compared via a two-way ANOVA (Fig. 5d, top). Additionally, the average total number of CK19^+^ surfaces for the non-fibrotic group and cirrhotic group were plotted and compared via an unpaired t-test (Fig. 5d, bottom).

## Statistics

Statistical comparisons were performed using a two-tailed Mann-Whitney test, a two-way ANOVA with a Bonferroni multiple comparisons test, or a two-tailed, unpaired t-test. Sample sizes for each dataset and relevant statistics are described in figure legends. All *p*-values <0.05 were considered statistically significant.

## Acknowledgments

This research was funded by NIH NIDDK (R01DK128551) (K.R.S.), the NIH Environmental Pathology and Toxicology Training Grant T32 ES007032 (W.B.F), the NCATS Translational Research Training Program TL1 TR002318 and the NIGMS Molecular Medicine Training Grant T32 GM095421 (C.L.F), and the NIH NIAAA R01AA026302 (R.M.C.). Any opinions, findings, conclusions, or recommendations expressed in this material are those of the author(s) and do not necessarily reflect the views of the funders. Schematics of liver tissue and liver lobules were created using the BioRender web application. We thank Dr. Dale Hailey from the UW Garvey Imaging Core for assistance with imaging. We thank all patient tissue donors for their generous contributions to science. We acknowledge that the experiments described in this text were conducted on the land of the past, present, and future Coast Salish peoples, the land that touches the shared waters of all tribes and bands within the Suquamish, Tulalip, and Muckleshoot nations; we honor the land itself and these Coast Salish tribes. The authors further acknowledge that papers authored by women and scholars from underrepresented racial and ethnic groups are systematically under cited and have made every attempt to reference relevant papers in a manner that is equitable in terms of gender and racial representation.

## Supplementary Figures

**Supplemental Figure 1:**
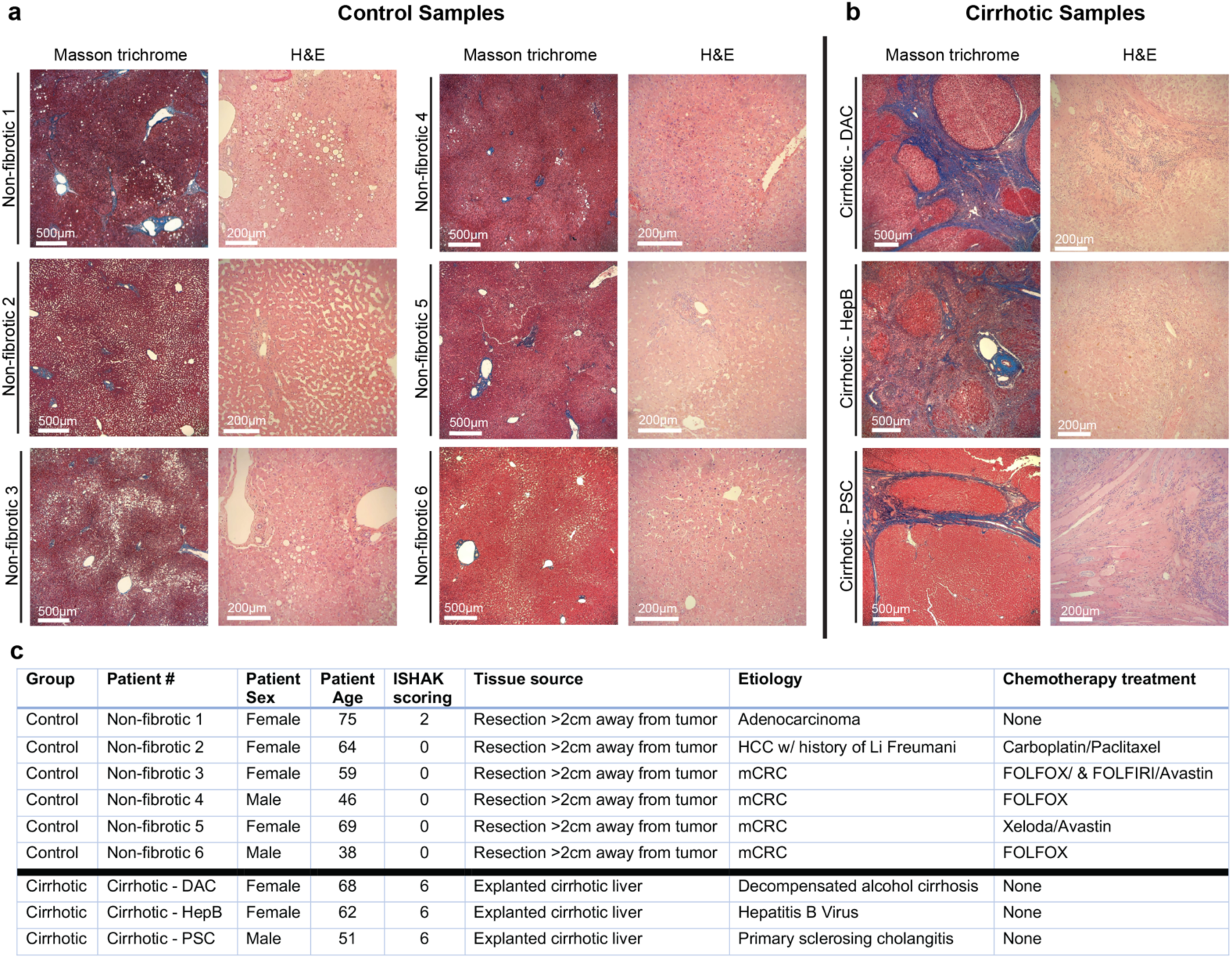
**a, b** Masson trichrome and hematoxylin and eosin (H&E) staining for control samples (**a**, n=6) and cirrhotic samples (**b**, n= 3; DAC, decompensated alcohol-associated cirrhosis; HepB, hepatitis B virus; PSC, primary sclerosing cholangitis). For Masson trichrome staining, collagen fibers stain blue, nuclei stain dark blue or black, and muscle fibers, cytoplasm, and keratin stain red. For the hematoxylin and eosin staining, nuclei stain blue via hematoxylin and extracellular matrix and cytoplasm stain red/pink via eosin. **c**, Patient information table showing group, patient biological sex, age, blinded pathologist ISHAK scoring, tissue source, etiology and chemotherapy treatment if applicable (HCC, hepatocellular carcinoma; mCRC, metastatic colorectal cancer; FOLFOX, folinic acid/fluorouracil/oxaliplatin; FOLFIRI, folinic acid/fluorouracil/irinotecan; Avastin, bevacizumab; Xeloda, capecitabine).

**Supplemental Figure 2:**
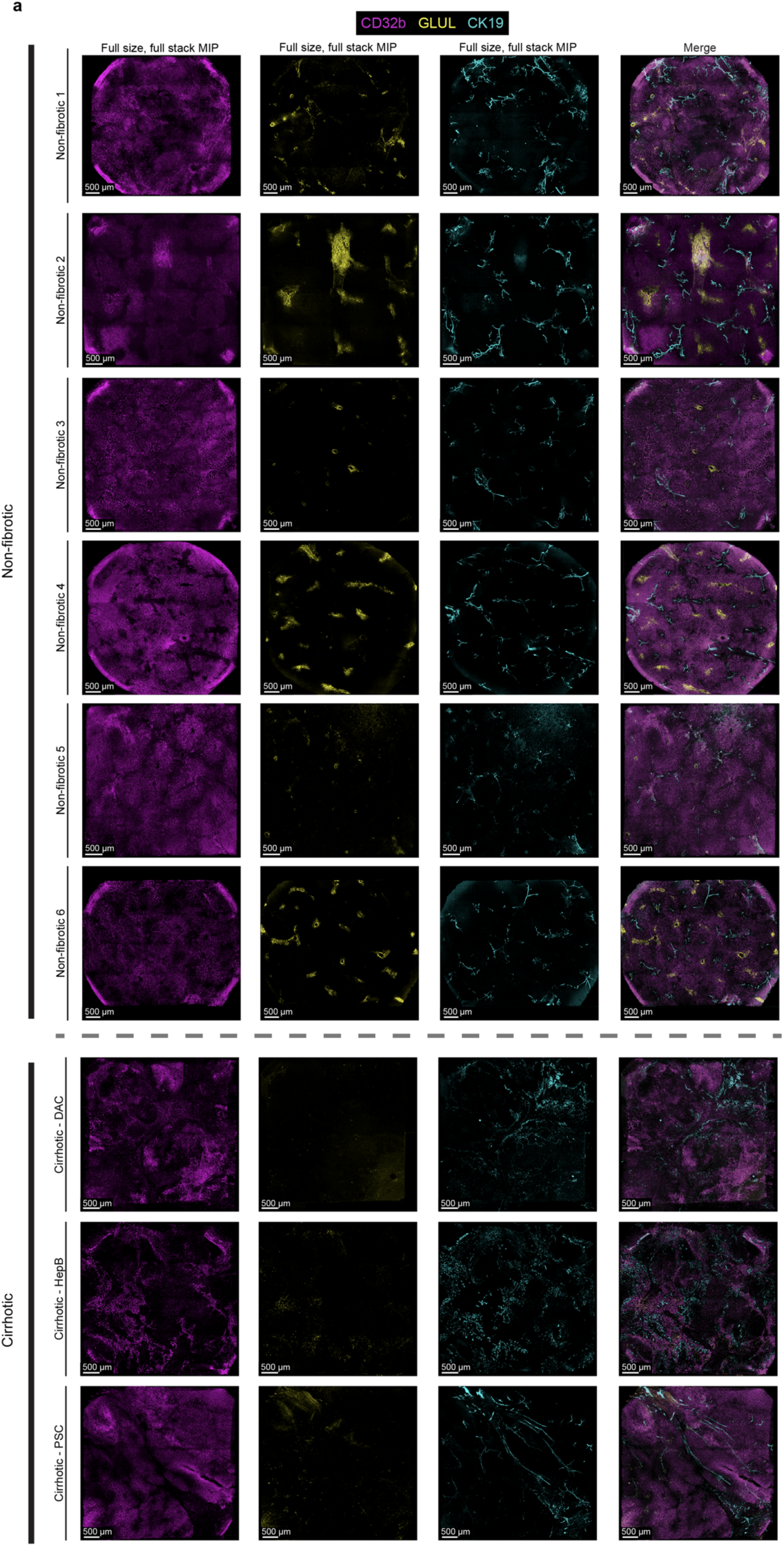
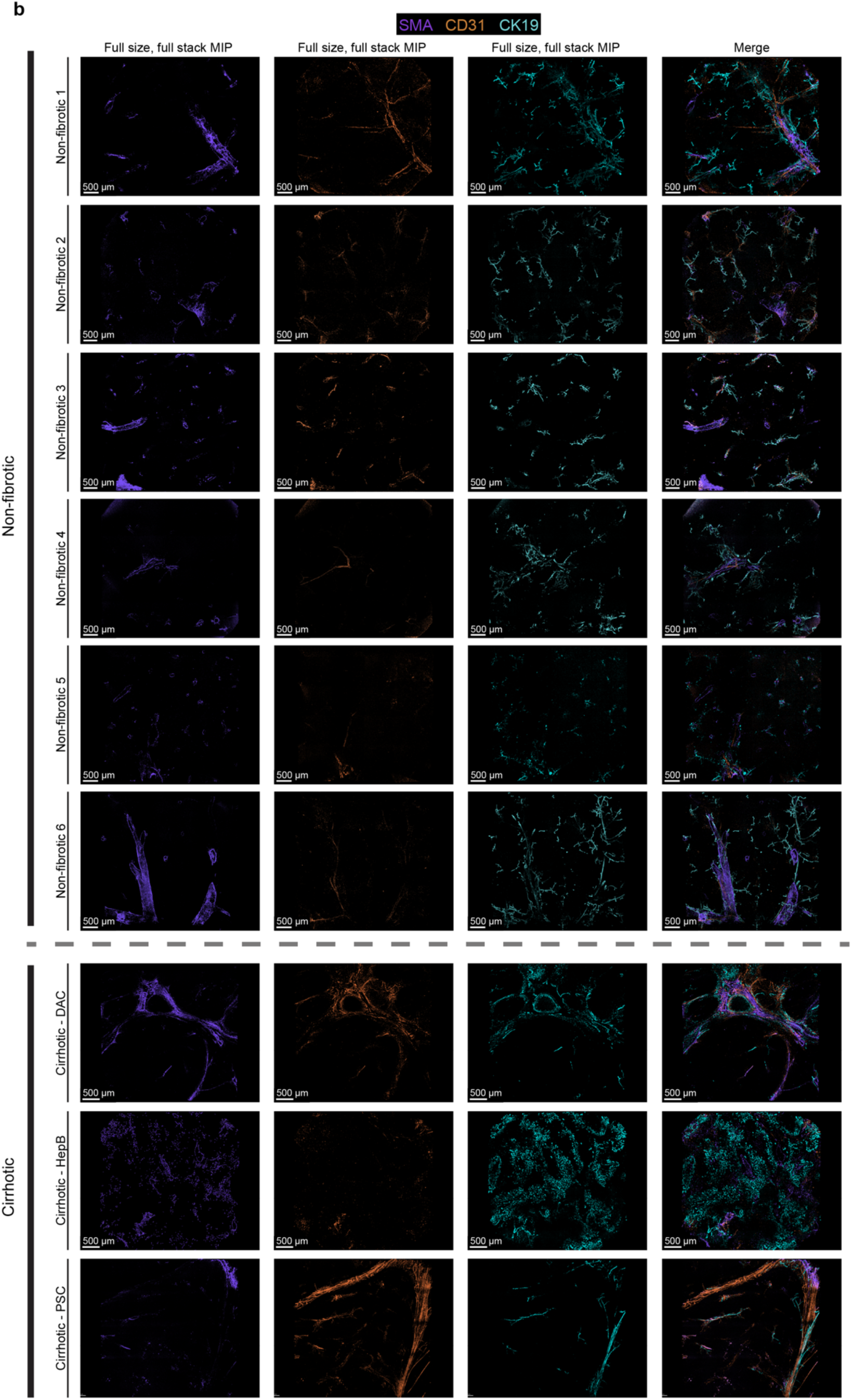
**a,b**, En face, full-stack MIP images (10x,16-tile merge) of immunofluorescence signals CD32b (magenta), glutamine synthetase (GLUL; yellow), and CK19 (cyan) (**a**) as well as smooth muscle actin (SMA; purple), CD31 (orange) and CK19 (cyan) (**b**) for non-fibrotic control tissues and cirrhotic tissues (DAC, decompensated alcohol-associated cirrhosis; HepB, hepatitis B virus; PSC, primary sclerosing cholangitis).

**Supplemental Figure 3:**
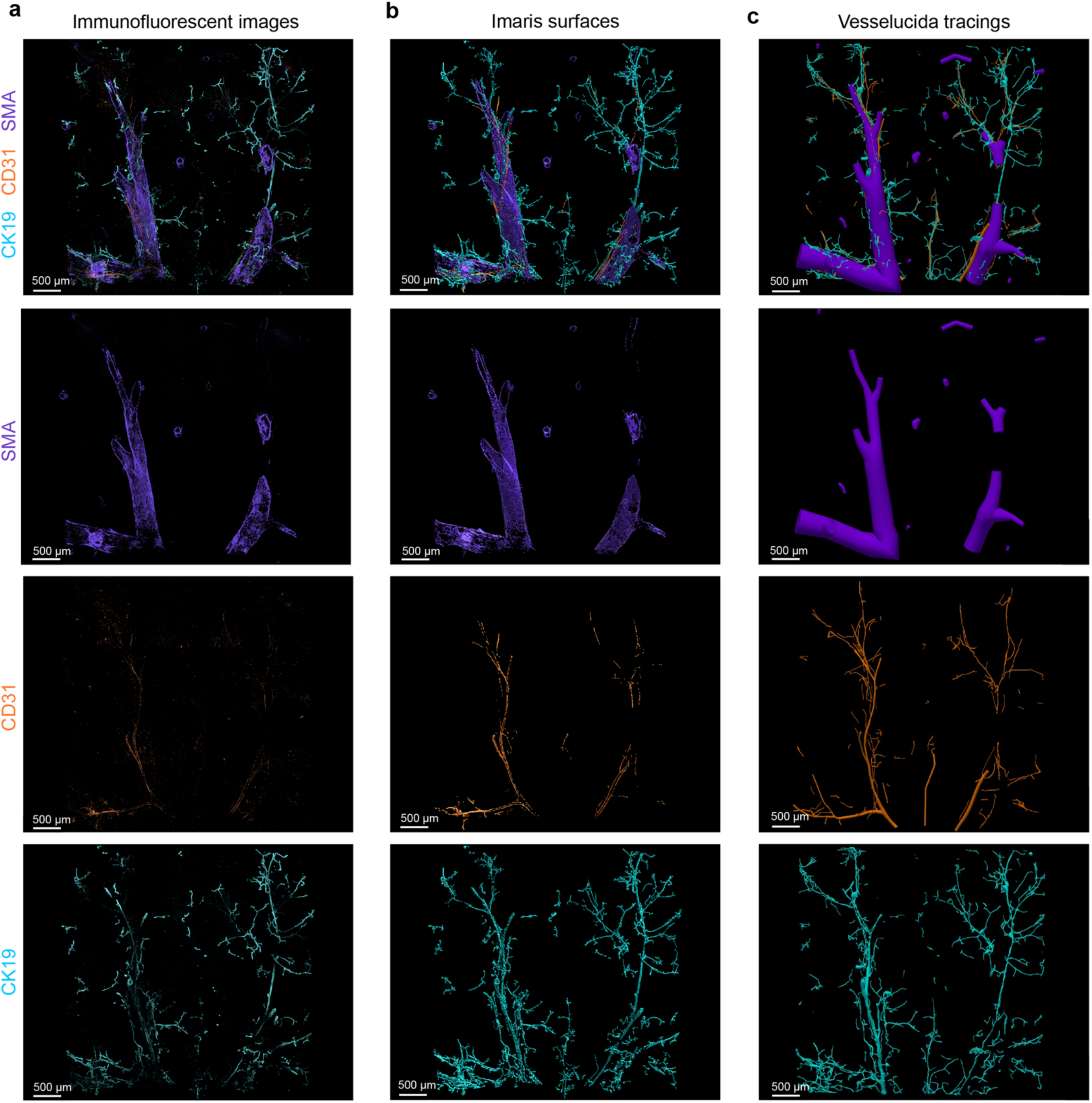
**a**, Immunofluorescent images of control tissue with SMA (purple) staining smooth muscle cells of central and portal veins, CD31 (orange) labeling hepatic arterial endothelial cells, and CK19 (cyan) marking biliary epithelial cells. **b**, Imaris surface renderings of immunofluorescent signals in **a**. **c**, Vesselucida tracings of immunofluorescent signals in **a**.

**Supplemental Figure 4:**
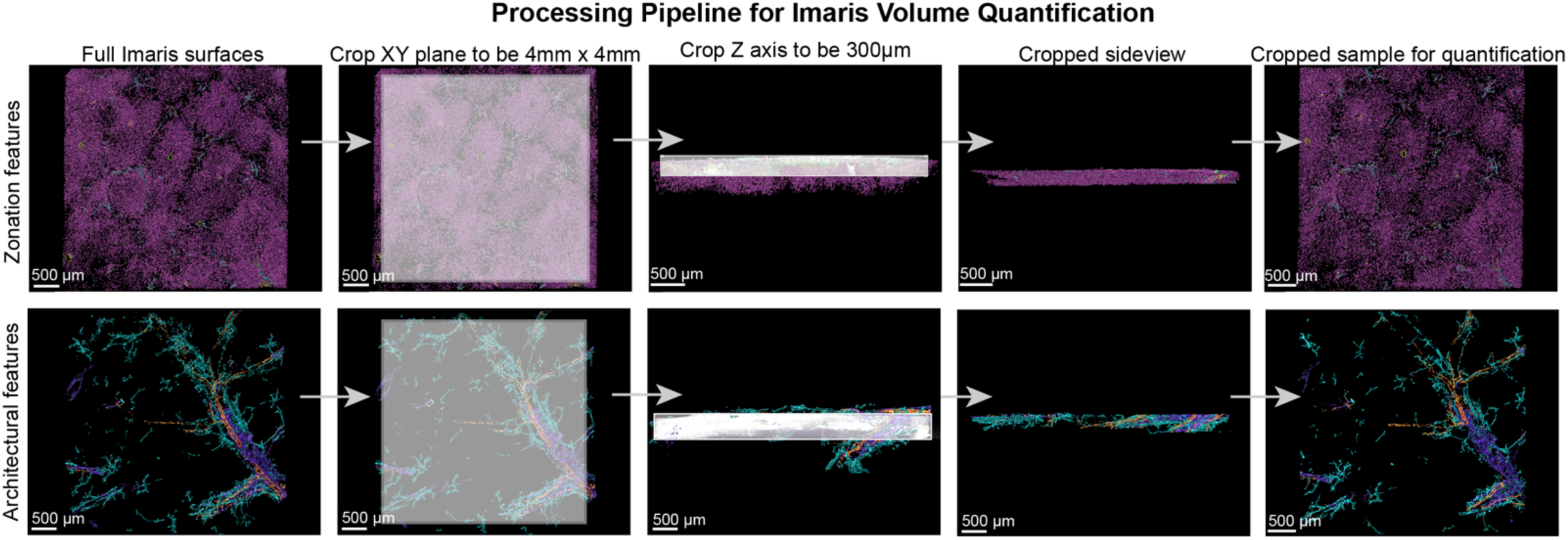
Imaris processing pipeline for visualization and quantification of zonation features (top) and architectural features (bottom).

**Supplemental Figure 5:**
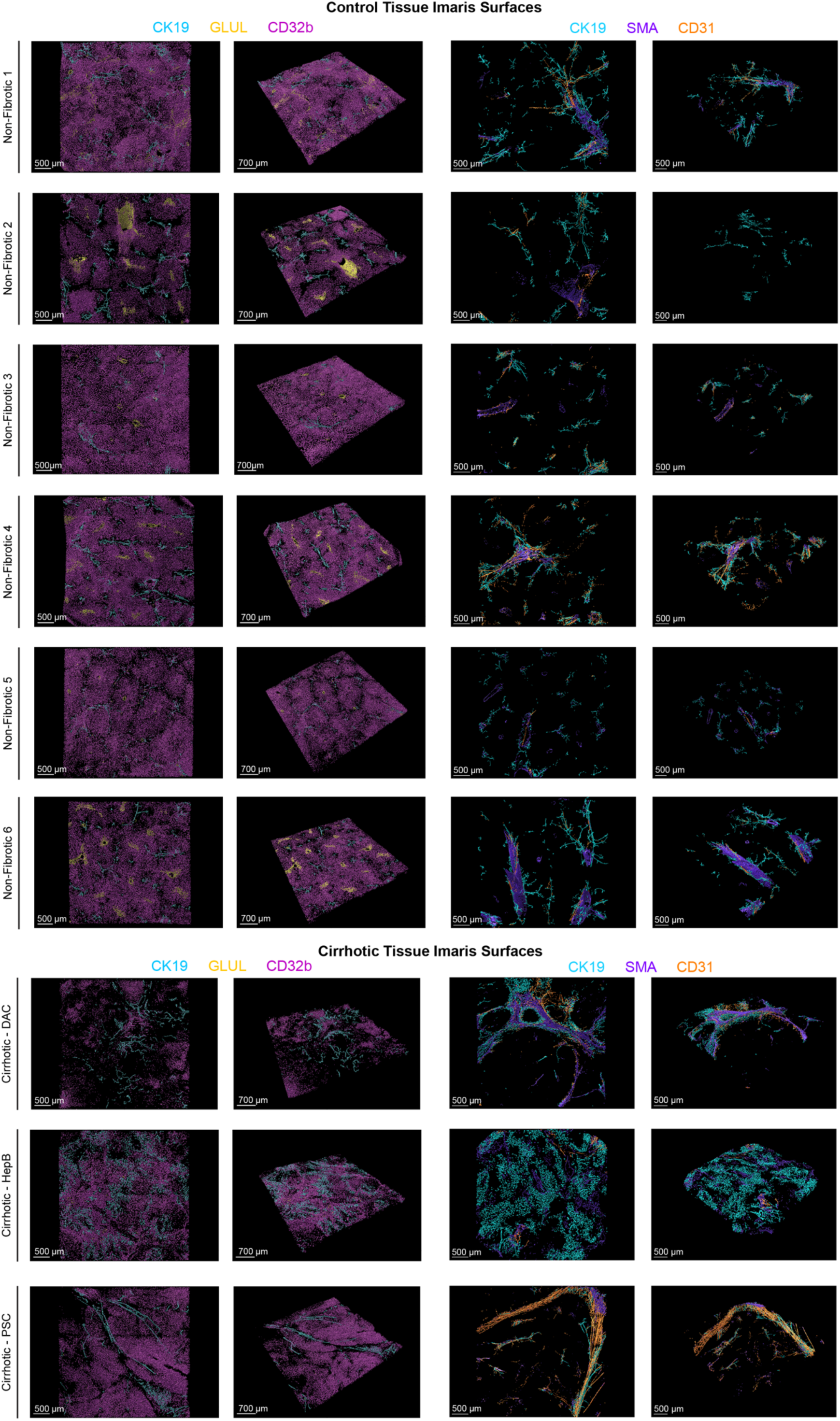
Depictions of Imaris surfaces reconstructing zonation features (left) and architectural features (right) for all patients from the non-fibrotic control group (n=6) and the cirrhotic group (n=3; DAC, decompensated alcohol-associated cirrhosis; HepB, hepatitis B virus; PSC, primary sclerosing cholangitis).

**Supplemental Figure 6:**
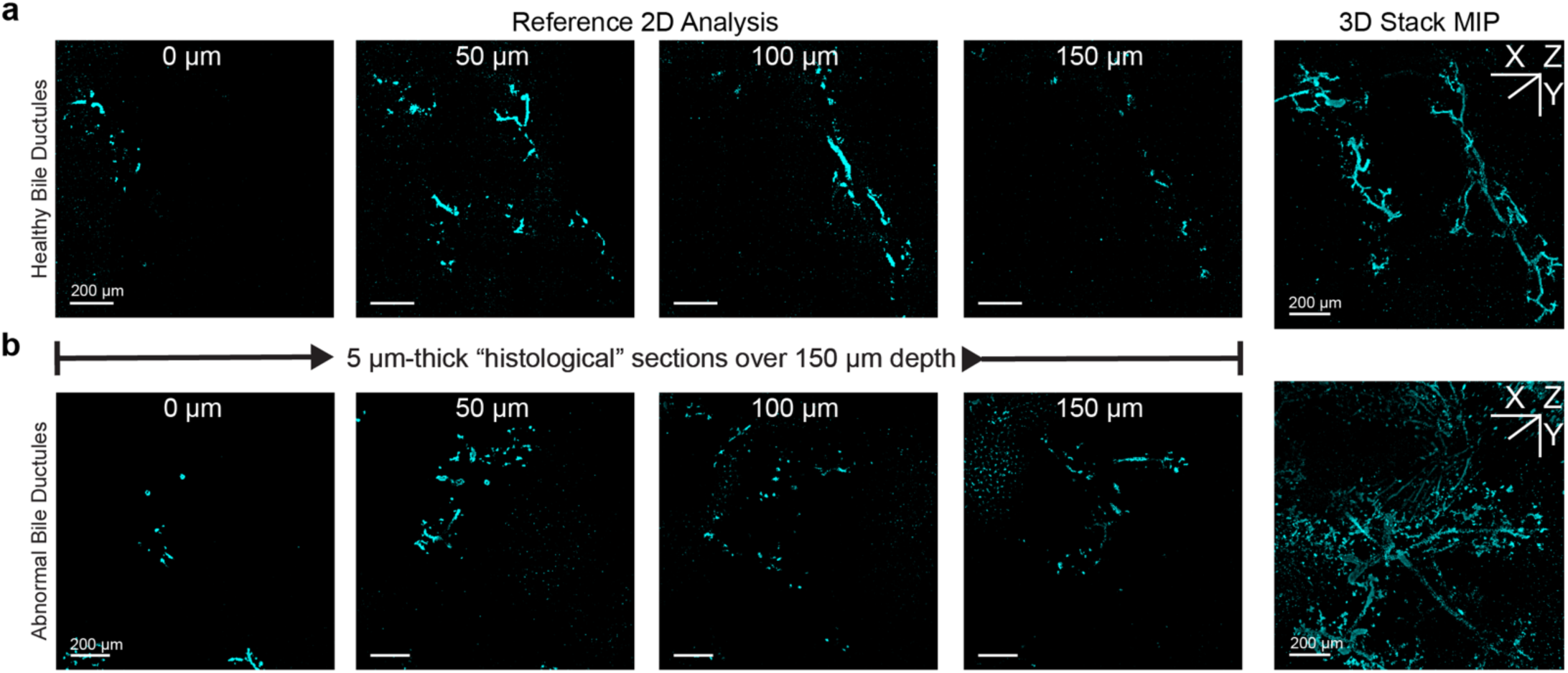
**a,b**, Left, Reference 2D histological analyses of 5µm-thick sections at various z-depths ranging 0 µm to 150 µm of healthy bile ducts (**a**) and abnormal bile ducts (**b**). Right, 3D Stack MIP of the same CK19^+^ networks.

## Author Contributions

**W.B.F.** developed and performed image processing pipelines; created 3D computational reconstructions of immunofluorescent signals using Imaris software; conceived and implemented quantitative measurements; and contributed to writing the manuscript. **C.L.F.** designed and performed immunostaining and tissue clearing strategies; imaged human liver samples; performed Vesselucida software reconstructions; conceived and implemented quantitative measurements; and contributed to writing the manuscript. **H.L.K.** collected patient information and performed fixation and primary tissue handling of human liver samples. **S.P.S.** assisted with staining, clearing, and imaging human liver samples as well as Vesselucida software reconstructions. **J.T.C** advised on imaging and clearing methodologies. **M.M.Y.** performed blinded ISHAK scoring as well as advised on pathological observations. **R.S.W.Y.** performed surgical resection of non-fibrotic human liver samples during partial hepatectomy and procured cirrhotic samples from excised human livers. **R.M.C.** advised on pathological findings presented in the manuscript. **K.R.S** guided scientific exploration as the primary advisor of trainees W.B.F., C.L.F., and S.P.S. for the duration of work and contributed to editing the manuscript.

